# White matter microglia morphological changes with aging in guinea pig offspring born growth restricted

**DOI:** 10.1101/2024.07.04.602101

**Authors:** Timothy B. Nunes, Karen L. Nygard, Marc C.J. Courchesne, Shawn N. Whitehead, Bryan S. Richardson, Timothy R.H. Regnault

## Abstract

Fetal growth restriction is implicated in the programming of later-life neurodegeneration. We hypothesized that growth restricted offspring would show accelerated changes to microglial white matter morphology, relative to controls.

Control guinea pig sows were fed *ad libitum*, while maternal nutrient restriction sows received 70% of control diet switched to 90% from mid-gestation. Offspring were sacrificed at ∼26 days (neonate) or ∼110 days (adult) postpartum. Coronal brain sections from the frontal cortex were subject to IBA1-staining for microglial detection and analyzed by machine learning software.

At birth, total body weight of growth restricted offspring was reduced relative to control (p<0.0001) with postnatal catch-up growth observed. Microglial density was reduced in the corpus callosum of control (p<0.05) and growth restricted (p=0.13) adults, relative to neonates. Adults from both groups showed greater IBA1-positive area in the cingulum and periventricular white matter (p<0.05) and increased microglial fractal dimension in the corpus callosum (p<0.10) and periventricular white matter (p<0.05), relative to neonates.

At the timepoints studied, we report age-related changes in white matter microglial morphology. However, maternal nutrient restriction leading to fetal growth restriction in guinea pigs does not appear to exacerbate these white matter microglia morphological changes as a marker for later-life neurodegeneration.

## Introduction

About 5-10% of all live human births are affected by fetal growth restriction (FGR) with maternal nutrient restriction (MNR) a major contributing factor (Fowden et al., 2006; Kamphof et al., 2022). In FGR offspring there is an increased risk of neurodegenerative pathologies in later life and more so in those showing postnatal catch-up growth (Fowden et al., 2006; Miller et al., 2016). Microglia-mediated activation and neuroinflammation appears to be a key driver of these neurological abnormalities in FGR offspring (Zinni et al., 2021). Grey matter microglial activation is traditionally associated with the presentation of dementia and Alzheimer’s disease, however recent findings implicate white matter pathology in earlier stages of neurodegeneration (Levit et al., 2019; Raj et al., 2017). Single-cell RNA sequencing has revealed that white matter microglia exhibit a transcriptome signature transitioning towards disease-associated microglia with aging (Safaiyan et al., 2021). These findings were reported in murine models of Alzheimer’s disease, and therefore characterization of white matter microglial morphology in FGR models is needed.

Ionized calcium-binding adaptor protein-1 (IBA1) is constitutively expressed by microglia and upregulated during activation to facilitate morphological changes observed (Hovens et al., 2014). As microglia transition to an inflammatory ameboid phenotype, cell processes retract and the soma grows. Thus, classical microglial activation may be characterized by an increased cell body to cell size ratio. To characterize subtle morphological changes, fractal dimension analysis provides an index of microglial branching complexity and number of primary processes (Karperien et al., 2013; Morrison et al., 2017).

We have previously shown the utility of guinea pigs in modelling MNR-induced FGR with postnatal catch-up growth (Nevin et al., 2018). Furthermore, non-transgenic guinea pigs closely mirror in-utero development and hallmarks of brain aging in humans (Wahl et al., 2022) and are an established model for studying the developmental origins of health and disease—including later-life Alzheimer’s (Sharman et al., 2013). Thus, we sought to determine whether MNR-induced FGR in guinea pig offspring would show accelerated changes to microglial morphology with aging. Microglial morphology of neonates and adults was examined for the corpus callosum (CC), cingulum (C), and periventricular white matter (PV). We assessed microglial density, percent area, classical activation, and fractal dimension in these white matter regions of interest. It was hypothesized that FGR would amplify age-related increases in these measures of microglial migration, enhanced phagocytosis, and primary process number.

## Methods

### Ethics Statement

Guinea pig brain tissue samples used in this experiment were provided from a previous study (Nevin et al., 2018), which employed an established model of moderate MNR in guinea pigs (Elias et al., 2016). Ethics approval for experimental procedures was obtained from Western University’s Animal Care Committee (Animal Use Protocol 2014-027). Continued ethics monitoring was performed in accordance with Western University Policy and Canadian Council on Animal Care guidelines.

### Animal feeding, pupping, and necropsies

Forty-one Dunkin-Hartley female guinea pigs (Charles River Laboratories, Sherbrooke, QC) aged 4-6 months were housed in individual enclosures, at a temperature of 25°C, with an automated 12-hour light-dark cycle. After two weeks of acclimation, sows were randomly assigned to a specific feeding protocol for a minimum of 4 weeks prior to mating with control diet boars. Control sows were fed *ad libitum* (Guinea Pig Diet 5025; LabDiet, St. Louis, MO), while MNR sows were fed 70% of what control sows consumed, determined as an average of food intake normalized to total body weight. At mid-pregnancy, from 35 days of gestation and onwards, MNR group food intake was increased to 90% of what control sows consumed. Notably, this dietary regime results in actual food consumption by the MNR animals of ∼65-70% of that of the control animals throughout pregnancy (Elias et al., 2016).

Pups were born spontaneously at a mean gestational age of 68 days, and birth weights were recorded within 24h of birth. Offspring from control sows with a birth weight greater than 95g were deemed appropriate for gestational age controls (Control n=29), while offspring from MNR sows with a birth weight below 85g were considered FGR (FGR-MNR n=26). These birth weight thresholds extrapolated from our fetal study using moderate MNR (Elias et al., 2016). After weaning, all pups received food *ad libitum*.

Twenty-seven neonates were randomly selected for necropsy at a mean of 26 days postnatal to provide the neonatal cohort: male neonate Control (n=7), female neonate Control (n=8), male neonate FGR-MNR (n=6), and female neonate FGR-MNR (n=6). The remaining offspring matured to adulthood and were necropsied at a mean of 110 days postnatal to provide the adult cohort: male adult Control (n=6), female adult Control (n=8), male adult FGR-MNR (n=6), and female adult FGR-MNR (n=8). Offspring were weighed prior to sacrifice and then euthanized with an intraperitoneal injection of 0.3mL pentobarbital sodium (Euthanyl; MTC Pharmaceuticals, Cambridge, ON). Necropsy weights of the brain and liver were recorded; additionally, these organs were fixed in 4% paraformaldehyde for 24 hours, rinsed in phosphate-buffered saline (PBS) three times at 2-hour intervals, and dehydrated in 70% ethanol for 2 weeks. Samples were then blocked in paraffin wax for future histochemical analyses.

### IBA1 Immunohistochemistry, Image Acquisition, and Analysis

A rotary microtome was used to cut 5μm coronal slices, corresponding to regions #720 to #760 of the *Cavia porcellus* Comparative Mammalian Brain Collection from the University of Wisconsin (Welker et al., 2010). Samples were mounted on 1.5mm Superfrost Plus Slides (VWR Scientific, Westchester, PA). All slides were stained on the same day with the same reagent pool to minimize variation in stain intensity.

Samples were deparaffinized with three consecutive 5-minute xylene washes, rehydrated in five progressive ethanol washes for 2 minutes each (100%, 100%, 90%, 90%, 70%), rinsed in running tap water for 5 minutes, and then submerged in reverse osmosis water for 1 minute. Heat-induced epitope retrieval was performed with slides submerged in 10mM sodium citrate at pH 6.0, using the 2100 Antigen Retriever (Aptum Biologics Ltd, Southampton, UK). Slides were gradually cooled to room temperature for 2 hours, rinsed with running tap water for 5 minutes, submerged in reverse osmosis water for 1 minute, and then rinsed with PBS. All PBS rinses were performed twice at 5 minutes intervals, at room temperature. Endogenous peroxidases were quenched with 3% hydrogen peroxide dissolved in PBS for 10 minutes. Slides were rinsed with PBS and then blocked with Background Sniper (BS966; Biocare Medical, Pacheco, CA) for 7 minutes. After a PBS rinse, samples were incubated with polyclonal rabbit anti-IBA1 primary antibody (1:3000, #019-19741, Wako Chemicals USA, VA) at 4°C overnight, in a covered humidity chamber. The primary antibody dilution was prepared with Dako antibody diluent (#S0809, Agilent Technologies Inc, Santa Clara, CA). One slide from each condition was incubated with diluent only and one slide from each condition was incubated with Dako rabbit non-immunized immunoglobulin fraction (#X0903, Agilent Technologies Inc, Santa Clara, CA) to serve as negative controls.

The next day, slides were rinsed with PBS and incubated for 40 minutes at room temperature with horseradish-peroxidase polymer horse anti-rabbit secondary antibody (MP-6401-15, Vector Laboratories, Burlington, ON) in a covered humidity chamber. Slides were then rinsed with PBS. To visualize bound antibody, samples were incubated with 3,3’-diaminobenzidine chromogen (#980681, MP Biomedicals, Santa Ana, CA) for 2 minutes and rinsed in running tap water for 5 minutes. Samples were counterstained in Mayer’s Hematoxylin for 70 seconds and immediately rinsed in running tap water for 5 minutes. Samples were dehydrated in five progressive ethanol washes for 2 minutes each (70%, 90%, 90%, 100%, 100%) and cleared with three consecutive 5-minute xylene washes. Slides were cover slipped with Permount (SP15-100, Fisher Scientific, Waltham, MA).

Image acquisition and analysis was performed by a researcher (T.N.) blinded to the experimental groups. Images were acquired for select brain regions of interest: the corpus callosum (CC), cingulum (C), and periventricular (PV) white matter. For each brain region, Z-stacks (9 planes over a depth of 4μm) for four different images were captured using an Eclipse Ti2-E inverted microscope (Nikon Instruments Inc, Melville, NY) with a 40x objective lens. For each sample the four images within each region were averaged as technical replicates. Z-stacks were then compiled into extended depth-of-focus TIFF files for analysis, to promote re-joining of complex 3D microglial processes with their associated cell bodies. All images were collected with identical illumination settings and an exposure time of 18ms to minimize variation in capturing.

Image analysis was performed with Image-Pro Premier 9.2 software (Media Cybernetics Inc, Rockville, MD). Regions of interest were drawn as needed for narrow regions, such as the corpus callosum. Thresholds were established to detect positive IBA1-staining and the same thresholds were applied uniformly across all images. Several images across the brain regions of interest were used to establish a machine learning algorithm with Image-Pro’s “smart selection” feature to sort IBA1-positive cell bodies with attached processes, IBA1-positive independent processes, and background (**Supp. Fig. 1**). The machine learning algorithm was then applied across all images with data collected for IBA1-positive cells per mm^2^, percent IBA1-positive stained area, cell body to cell size ratio, and fractal dimension. Cell density and fractal dimension were assessed using IBA1-positive cell bodies with attached processes. Cell body to cell size ratio was calculated on each image as the sum of pixels from IBA1-positive cell bodies divided by the sum of pixels from IBA1-positive cell bodies and IBA1-positive independent processes.

**Fig. 1.**
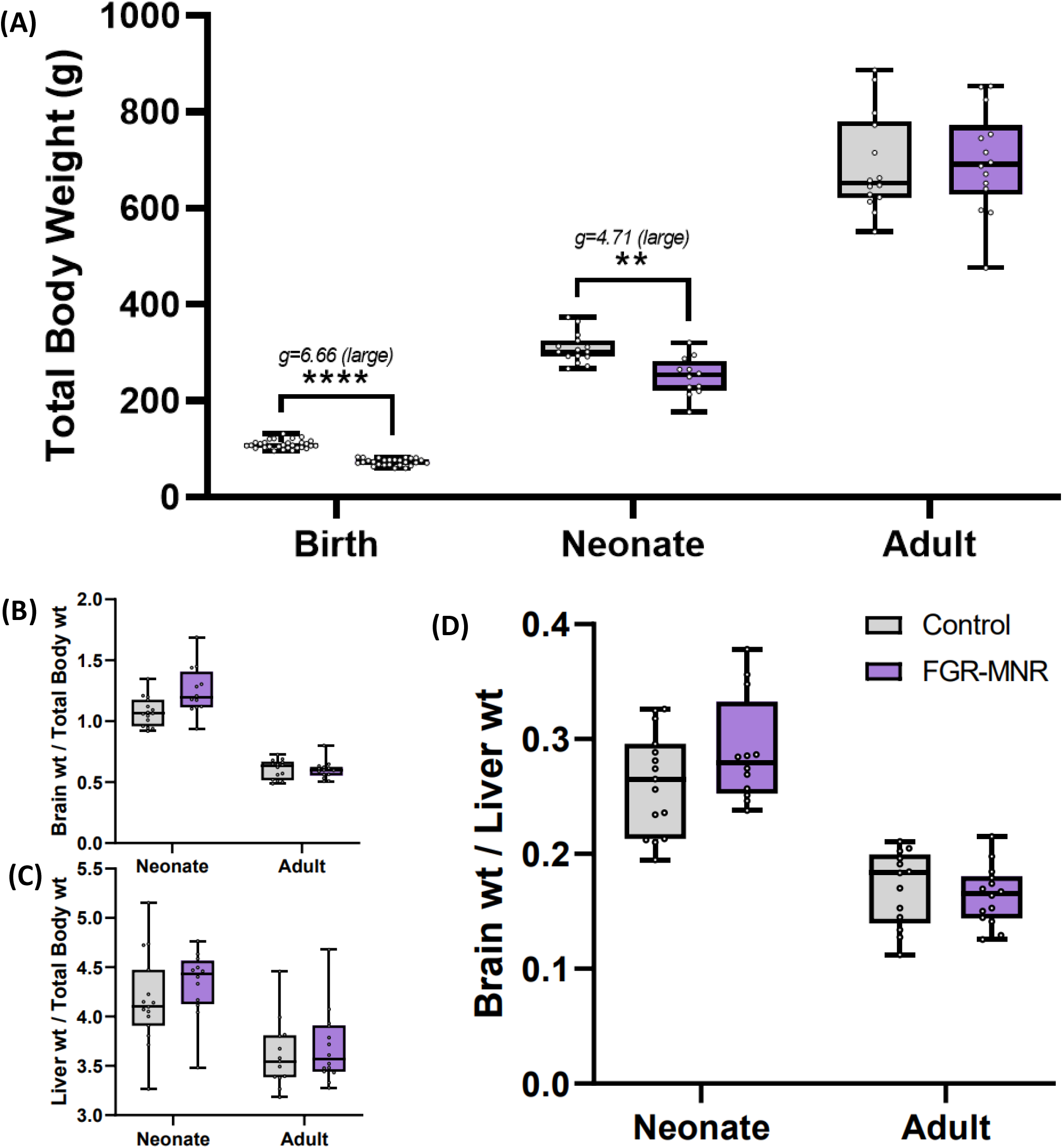
Maternal nutrient restriction produces asymmetrically growth restricted offspring, which exhibit postnatal catch-up growth. Total body weight **(A)**, normalized brain weight **(B)**, normalized liver weight **(C)**, and brain-liver ratios **(D)** for control (grey) and maternal nutrient restriction (purple) guinea pig offspring. Data are presented as median, 1st and 3rd quartiles, and extrema. (**p<0.01, ****p<0.0001; Hedge’s *g* effect sizes).

### Statistical Analysis

The ARRIVE guidelines stipulate the importance of addressing clustering effects in litter-based data (Sert et al., 2020). To appropriately assess variation between and within litters, we created linear mixed models and conducted estimated marginal means (emmeans) pairwise analysis—with litter identification codes assigned to each offspring as the random effect. These models were used to analyze differences across fixed effects of maternal diet and postnatal age. Sex was also included in the models, but the sample size was insufficient to report differences across sex. Statistical analyses were conducted using R 4.3.1 with several packages: *lme4* to construct the models, *lmerTest* to conduct Type III ANOVA with Satterhwaite’s method, and *emmeans* to conduct Tukey’s post-hoc pairwise analyses. Hedges’ *g* effect size analyses and correction for small-sample bias were conducted with the eff_size function in the *emmeans* package (Hedges, 1981). GraphPad Prism 9.5.1 was used to generate graphics. Results are reported as group emmeans ± standard error of the mean. For all analyses, p<0.05 with *g*>1 was considered significant.

## Results

### Growth characterization

FGR-MNR total body weights were decreased 32% at birth compared with those of controls (73.60±1.80g vs 109.46±1.58g, p<0.001), but were only decreased 17% in the neonatal groups at postnatal day 26 (256.71±11.88g vs 307.77±10.06g, p<0.01), and with no difference in the adult groups at postnatal day 110 (688.89±27.06g vs 688.95±24.51g, p=0.99) (**Fig. 1A**). Brain and liver weights of offspring were recorded at necropsy and normalized to total body weight. FGR-MNR neonates (1.16±0.05) exhibited marginally greater normalized brain weight, relative to control neonates (1.14±0.05) (p=0.61), but catch-up growth had occurred by adulthood (p=0.95) (**Fig. 1B**). No differences in normalized liver weight were observed in neonates (p=0.72) or adults (p=0.96) (**Fig. 1C**). Brain to liver ratios were calculated based on organ weights at necropsy. While not significant, FGR-MNR neonates (0.29±0.01) exhibited a marginally greater brain to liver ratio, relative to control neonates (0.26±0.01) (p=0.50), with this relationship diminishing by adulthood (p=0.7840) (**Fig. 1D**).

### IBA1-positive cell count

Total IBA1-positive cell count per mm^2^ was collected for the CC, C, and PV, as a proxy of microglial density. In the CC, control adults exhibited a reduced IBA1-positive cell count relative to control neonates (130.28±12.19cells/mm^2^ vs 177.42±12.40cells/mm^2^, p<0.05). FGR-MNR adults also exhibited a reduced IBA1-positive cell count relative to FGR-MNR neonates in the CC (129.80±13.07cells/mm^2^ vs 172.82±13.67cells/mm^2^, p=0.13, *large g*=1.15), although not significant. No differences in IBA1-positive cell count were observed across maternal diet or postnatal age groupings in the C and PV. (**Fig. 2A**).

**Fig. 2.**
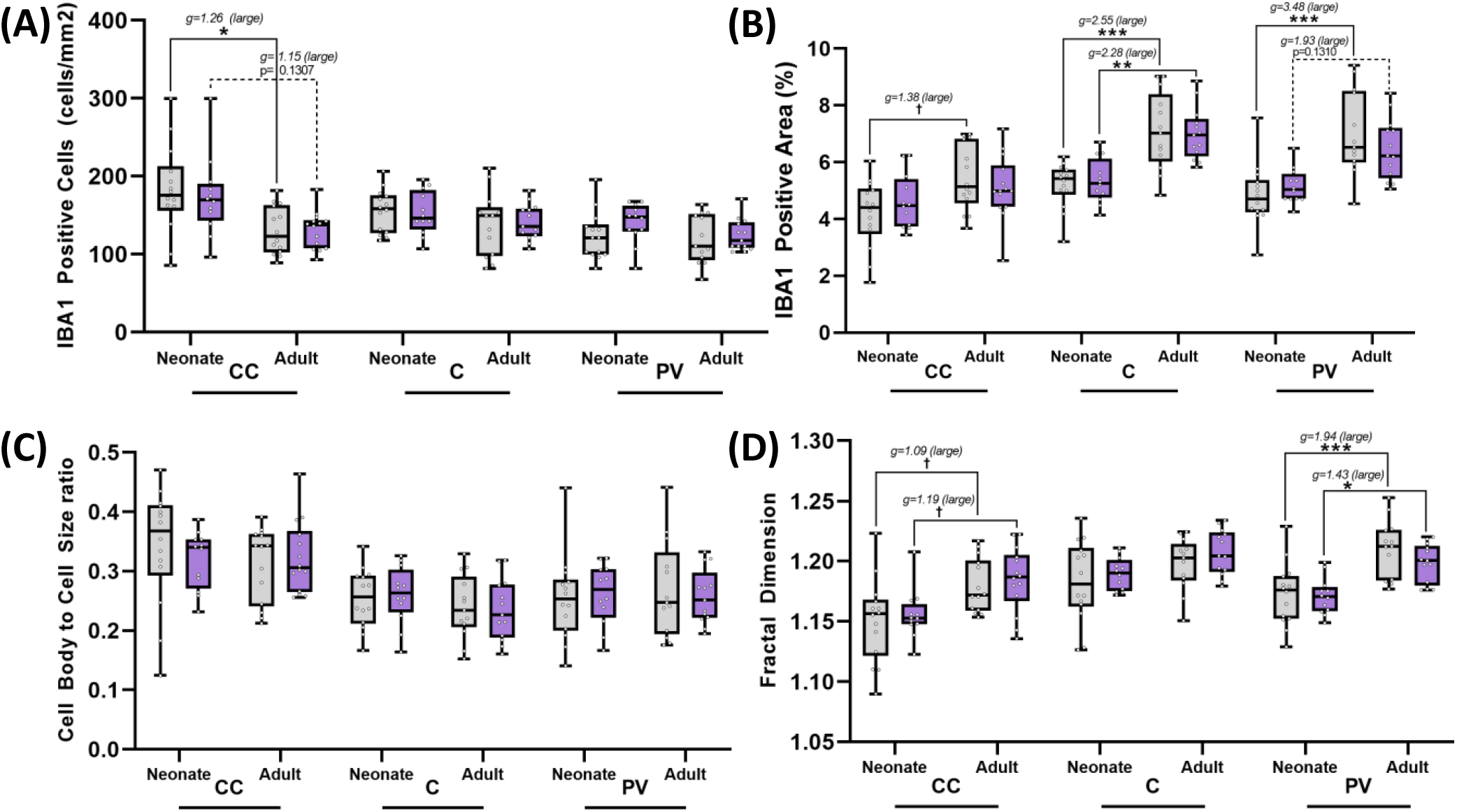
Microglial morphology in the corpus callosum, cingulum, and periventricular white matter is altered with aging, but is not exacerbated by fetal growth restriction. Coronal brain sections were subject to IBA1-staining for microglial detection and analyzed by machine learning software. Normalized IBA1-positive cell count **(A)**, percent IBA1-positive area **(B)**, microglia cell body to cell size ratio **(C)**, and microglia fractal dimension **(D)** for control (grey) and maternal nutrient restriction (purple) guinea pig offspring. Data are presented as median, 1st and 3rd quartiles, and extrema. CC = corpus callosum. C = cingulum. PV = periventricular white matter. (†p<0.1, *p<0.05, **p<0.01, ***p<0.0001; Hedge’s *g* effect sizes).

### Percent IBA1-positive stained area

Percent IBA1-positive stained area was collected for the CC, C, and PV, as a proxy of total microglial volume. Control adults exhibited increased percent IBA1-positive area relative to control neonates in the CC (5.37±0.32% vs 4.17±0.32%, p<0.1, *large g*=1.38), the C (7.14±0.30% vs 5.26±0.30%, p<0.001), and the PV (7.04±0.34% vs 4.80±0.34% p<0.001). Similarly, FGR-MNR adults exhibited increased percent IBA1-positive area relative to FGR-MNR neonates in the C (7.02±0.34% vs 5.33±0.34%, p<0.01) and trended towards this increase in the PV (6.58±0.39% vs 5.34±0.39%, p=0.13, *large g*=1.93). No differences in IBA1-positive area were observed between control and FGR-MNR cohorts at either age point in the white matter regions analyzed. (**Fig. 2B**).

### Microglial cell body to cell size ratio

Microglial cell body to cell size ratio was collected for the CC, C, and PV, as an indicator of classical microglial activation. No differences in this ratio were observed within or between control and FGR-MNR cohorts at either age point studied. (**Fig. 2C**).

### Microglial fractal dimension

Microglia fractal dimension was collected for the CC, C, and PV, as an index of branching complexity and total process number. In the CC, control adults exhibited increased fractal dimension relative to control neonates (1.18±0.01% vs 1.15±0.01%, p<0.1, *large g*=1.09), as did the FGR-MNR adults relative to the FGR-MNR neonates (1.18±0.01% vs 1.15±0.01%, p<0.1, *large g*=1.19). Similarly in the PV, control adults exhibited increased fractal dimension relative to control neonates (1.21±0.01% vs 1.17±0.01%, p<0.001), as did the FGR-MNR adults relative to the FGR-MNR neonates (1.20±0.01% vs 1.17±0.01%, p<0.05). No differences in fractal dimension were observed between control and FGR-MNR cohorts at either age point in the white matter regions analyzed. (**Fig. 2D**).

## Discussion

This study sought to determine whether FGR exacerbates microglial morphological changes with aging in white matter regions of interest. Our findings highlight age-related changes in white matter microglial morphology associated with neuroinflammation and neurodegeneration. In the corpus callosum, we report a significant increase in branching complexity and a reduction in microglial density with aging. In the cingulum and periventricular white matter, we have shown significantly increased IBA1-positive percent area with aging, as a proxy of total microglial volume. At the time points examined, FGR by total caloric MNR does not appear to exacerbate microglial morphological changes with aging in the corpus callosum, cingulum, or periventricular white matter.

To demonstrate the translatability of our guinea pig model to human FGR-MNR pregnancies, we examined growth characterization data of offspring at birth, neonate, and young adult stages of life. In human pregnancies, maternal malnourishment commonly results in asymmetrically growth restricted newborns (Sharma et al., 2016). Our growth characterization trends are consistent with previous reports that FGR by MNR in guinea pigs produces asymmetrical growth restriction with postnatal catch-up growth—representative of maternal malnourishment in humans (Elias et al., 2016; Kind et al., 2005). FGR-MNR offspring were born with significantly reduced total body weight, relative to control. Our group has previously shown significant increases in brain-liver ratios of FGR-MNR guinea pig fetuses at term, lending support to our asymmetrical growth restriction model (Maki et al., 2019). By postnatal day 26, FGR-MNR neonates were still significantly lighter and trended towards larger normalized brain weights compared to controls. This trend captures asymmetrical brain-sparing known to occur in malnourished human fetuses (Sharma et al., 2016). However, rapid catch-up growth was evident, as FGR-MNR neonates exhibited normalized liver weights within control levels. By postnatal day 110, no differences remained in organ weights or total body weights between FGR-MNR and control adults. The rapid postnatal catch-up growth we observed in FGR-MNR offspring is a known risk factor in later-life neurodegenerative disease (Fowden et al., 2006; Miller et al., 2016).

The main finding of our study was considerable changes in microglial morphology across the corpus callosum, cingulum, and periventricular white matter with aging. Microglial density was significantly reduced in the corpus callosum of controls adults compared to neonates, and this trend was also observed in FGR-MNR adults. The reduction in microglia within the corpus callosum is likely due to increased microglial migration with aging. Microglia have been shown to migrate via the corpus callosum towards spontaneous cortical microinfarcts associated with neurodegeneration (Lubart et al., 2021). We also report increased percent IBA1-positive area in the cingulum and periventricular white matter with aging, as a proxy of total microglial volume (Kongsui et al., 2014). Interestingly, this increase occurred with no significant changes in microglial cell body to cell size ratio. Thus, a uniform increase in microglial volume occurred, with the cell body and processes growing proportionally. This suggests microglia increased in size without fully transitioning to a classical ameboid inflammatory phenotype (Hovens et al., 2014). Our finding is supported by previous studies reporting enhanced phagocytic activity of microglia with aging in white matter tracts (Raj et al., 2017; Shobin et al., 2017). Lastly, we report increased fractal dimension in the corpus callosum and periventricular white matter with aging. While the number of processes leaving the microglia soma increases with age, the complexity of branching may be unaffected (Karperien et al., 2013). Therefore, our observed age-related increase in microglial fractal dimension can likely be attributed to an increase in primary processes.

While previous animal studies have reported white matter microglia-mediated neuroinflammation in FGR offspring, differences in experimental design may explain our results.

With a porcine model of spontaneous FGR, microglia-mediated inflammation has been reported in the periventricular white matter at postnatal days 1 and 4 (Wixey et al., 2019). Due to the remarkable plasticity of the perinatal brain, it is possible that acute neuroinflammatory changes present at birth could no longer be captured in microglial morphology at postnatal day 26 in our guinea pig model. For example, upregulation of proinflammatory cytokines in the white matter— such as tumor necrosis factor alpha, interleukin 1 beta, and interleukin 6—exhibited a marked decline by postnatal day 4 in FGR piglets (Wixey et al., 2019). With a rat model of FGR by uterine artery ligation, temporary changes in white matter microglia morphology have also been noted (Olivier et al., 2007). In the cingulum, higher numbers of ameboid microglia were reported in FGR offspring up to postnatal day 14, but this phenomenon diminished by postnatal day 21 (Olivier et al., 2007). When comparing these outcomes to our study, it is important to consider differences in CNS development timing between postnatal developers—such as rats—and prenatal developers— such as guinea pigs and humans (Morrison et al., 2018). Nonetheless, this rat study could indicate that ephemeral postnatal changes in microglial morphology may have been missed by the timing of our neonatal guinea pig collection.

Our study has further elucidated white matter microglial morphological changes with aging using modern machine learning analysis techniques. Specifically, our findings highlight age-related increases in microglia migration, enhanced phagocytosis, and primary process number in white matter regions of interest. The scope of our study was limited to select white matter regions in neonates and young adults, and we speculate that microglial white matter pathologies arising from FGR programming may only be observable perinatally and later in life (Gauvrit et al., 2022; Olivier et al., 2007; Wixey et al., 2019). At the time points studied, we have demonstrated that FGR by MNR in guinea pigs—a prenatal developer—does not accelerate changes to microglia morphology in the corpus callosum, cingulum, or periventricular white matter.

## Author Statements

## Acknowledgements

The authors thank Dr. Lannette Friesen-Waldner and Mr. Brian Sutherland for their assistance with animal support services.

## Competing interests

The authors declare there are no competing interests.

### Author Contributions (CRediT roles)

Conceptualization: BR & TR

Data curation: TN

Formal Analysis: TN

Funding acquisition: BR & TR

Investigation: TN, MC, & KN

Methodology: TN, MC, KN, & SW

Project administration: TR & BR

Resources: BR

Software: KN & TN

Supervision: BR & TR

Validation BR, KN, & TR

Visualization TN

Writing – original draft TN

Writing – review & editing TR, KN, & BR

### Funding

This research was supported by the London Health Sciences Centre Women’s Development Council and the NeuroDevNet Centre of Excellence.

### Data availability

Data generated or analyzed during this study are provided within the published article and its supplementary material.

## Supplementary Material

**Supp. Fig. 1.**
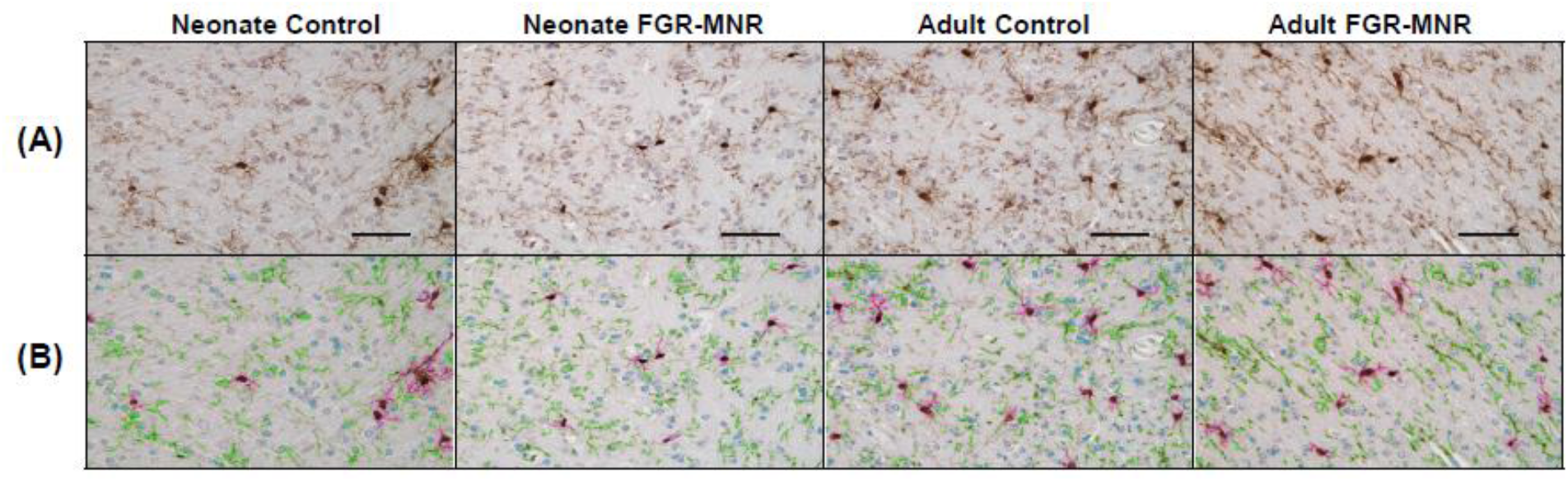
Machine learning sorting of microglial features. Coronal brain sections were subject to IBA1-staining for microglial detection and analyzed with ImagePro software. Thresholds were established to detect positive IBA1-staining and a machine learning algorithm was applied prior to quantification to sort microglial cell bodies with attached processes (pink), independent microglial processes (green), and background nuclei (cyan). Sorting methods were applied uniformly across all brain regions analyzed, with representative raw **(A)** and sorted **(B)** cingulum micrographs shown. Scale bar = 100μm.

## References

Elias, A.A., Ghaly, A., Matushewski, B., Regnault, T.R.H., Richardson, B.S., 2016. Maternal Nutrient Restriction in Guinea Pigs as an Animal Model for Inducing Fetal Growth Restriction. Reprod. Sci. 23, 219–227. 10.1177/1933719115602773

Fowden, A.L., Giussani, D.A., Forhead, A.J., 2006. Intrauterine Programming of Physiological Systems: Causes and Consequences. Physiology 21, 29–37. 10.1152/physiol.00050.2005

Gauvrit, T., Benderradji, H., Buée, L., Blum, D., Vieau, D., 2022. Early-Life Environment Influence on Late-Onset Alzheimer’s Disease. Front. Cell Dev. Biol. 10, 834661. 10.3389/fcell.2022.834661

Hedges, L.V., 1981. Distribution Theory for Glass’s Estimator of Effect size and Related Estimators. J. Educ. Stat. 6, 107–128. 10.3102/10769986006002107

Hovens, I., Nyakas, C., Schoemaker, R., 2014. A novel method for evaluating microglial activation using ionized calcium-binding adaptor protein-1 staining: cell body to cell size ratio. Neuroimmunol. Neuroinflammation 1, 82. 10.4103/2347-8659.139719

Kamphof, H.D., Posthuma, S., Gordijn, S.J., Ganzevoort, W., 2022. Fetal Growth Restriction: Mechanisms, Epidemiology, and Management. Matern.-Fetal Med. 4, 186. 10.1097/FM9.0000000000000161

Karperien, A., Ahammer, H., Jelinek, H.F., 2013. Quantitating the subtleties of microglial morphology with fractal analysis. Front. Cell. Neurosci. 7, 3. 10.3389/fncel.2013.00003

Kind, K.L., Roberts, C.T., Sohlstrom, A.I., Katsman, A., Clifton, P.M., Robinson, J.S., Owens, J.A., 2005. Chronic maternal feed restriction impairs growth but increases adiposity of the fetal guinea pig. Am. J. Physiol. Regul. Integr. Comp. Physiol. 288, R119–126. 10.1152/ajpregu.00360.2004

Kongsui, R., Beynon, S.B., Johnson, S.J., Walker, F.R., 2014. Quantitative assessment of microglial morphology and density reveals remarkable consistency in the distribution and morphology of cells within the healthy prefrontal cortex of the rat. J. Neuroinflammation 11, 182. 10.1186/s12974-014-0182-7

Levit, A., Regis, A.M., Gibson, A., Hough, O.H., Maheshwari, S., Agca, Y., Agca, C., Hachinski, V., Allman, B.L., Whitehead, S.N., 2019. Impaired behavioural flexibility related to white matter microgliosis in the TgAPP21 rat model of Alzheimer disease. Brain. Behav. Immun. 80, 25–34. 10.1016/j.bbi.2019.02.013

Lubart, A., Benbenishty, A., Har-Gil, H., Laufer, H., Gdalyahu, A., Assaf, Y., Blinder, P., 2021. Single Cortical Microinfarcts Lead to Widespread Microglia/Macrophage Migration Along the White Matter. Cereb. Cortex N. Y. N 1991 31, 248–266. 10.1093/cercor/bhaa223

Maki, Y., Nygard, K., Hammond, R.R., Regnault, T.R.H., Richardson, B.S., 2019. Maternal Undernourishment in Guinea Pigs Leads to Fetal Growth Restriction with Increased Hypoxic Cells and Oxidative Stress in the Brain. Dev. Neurosci. 41, 290–299. 10.1159/000506939

Miller, S.L., Huppi, P.S., Mallard, C., 2016. The consequences of fetal growth restriction on brain structure and neurodevelopmental outcome. J. Physiol. 594, 807–823. 10.1113/JP271402

Morrison, H., Young, K., Qureshi, M., Rowe, R.K., Lifshitz, J., 2017. Quantitative microglia analyses reveal diverse morphologic responses in the rat cortex after diffuse brain injury. Sci. Rep. 7, 13211. 10.1038/s41598-017-13581-z

Morrison, J.L., Botting, K.J., Darby, J.R.T., David, A.L., Dyson, R.M., Gatford, K.L., Gray, C., Herrera, E.A., Hirst, J.J., Kim, B., Kind, K.L., Krause, B.J., Matthews, S.G., Palliser, H.K., Regnault, T.R.H., Richardson, B.S., Sasaki, A., Thompson, L.P., Berry, M.J., 2018. Guinea pig models for translation of the developmental origins of health and disease hypothesis into the clinic. J. Physiol. 596, 5535–5569. 10.1113/JP274948

Nevin, C.L., Formosa, E., Maki, Y., Matushewski, B., Regnault, T.R.H., Richardson, B.S., 2018. Maternal nutrient restriction in guinea pigs as an animal model for studying growth-restricted offspring with postnatal catch-up growth. Am. J. Physiol.-Regul. Integr. Comp. Physiol. 314, R647–R654. 10.1152/ajpregu.00317.2017

Olivier, P., Baud, O., Bouslama, M., Evrard, P., Gressens, P., Verney, C., 2007. Moderate growth restriction: deleterious and protective effects on white matter damage. Neurobiol. Dis. 26, 253– 263. 10.1016/j.nbd.2007.01.001

Raj, D., Yin, Z., Breur, M., Doorduin, J., Holtman, I.R., Olah, M., Mantingh-Otter, I.J., Van Dam, D., De Deyn, P.P., den Dunnen, W., Eggen, B.J.L., Amor, S., Boddeke, E., 2017. Increased White Matter Inflammation in Aging- and Alzheimer’s Disease Brain. Front. Mol. Neurosci. 10, 206. 10.3389/fnmol.2017.00206

Safaiyan, S., Besson-Girard, S., Kaya, T., Cantuti-Castelvetri, L., Liu, L., Ji, H., Schifferer, M., Gouna, G., Usifo, F., Kannaiyan, N., Fitzner, D., Xiang, X., Rossner, M.J., Brendel, M., Gokce, O., Simons, M., 2021. White matter aging drives microglial diversity. Neuron 109, 1100-1117.e10. 10.1016/j.neuron.2021.01.027

Sert, N.P. du, Ahluwalia, A., Alam, S., Avey, M.T., Baker, M., Browne, W.J., Clark, A., Cuthill, I.C., Dirnagl, U., Emerson, M., Garner, P., Holgate, S.T., Howells, D.W., Hurst, V., Karp, N.A., Lazic, S.E., Lidster, K., MacCallum, C.J., Macleod, M., Pearl, E.J., Petersen, O.H., Rawle, F., Reynolds, P., Rooney, K., Sena, E.S., Silberberg, S.D., Steckler, T., Würbel, H., 2020. Reporting animal research: Explanation and elaboration for the ARRIVE guidelines 2.0. PLOS Biol. 18, e3000411. 10.1371/journal.pbio.3000411

Sharma, D., Shastri, S., Sharma, P., 2016. Intrauterine Growth Restriction: Antenatal and Postnatal Aspects. Clin. Med. Insights Pediatr. 10, 67–83. 10.4137/CMPed.S40070

Sharman, M.J., Moussavi Nik, S.H., Chen, M.M., Ong, D., Wijaya, L., Laws, S.M., Taddei, K., Newman, M., Lardelli, M., Martins, R.N., Verdile, G., 2013. The Guinea Pig as a Model for Sporadic Alzheimer’s Disease (AD): The Impact of Cholesterol Intake on Expression of AD-Related Genes. PloS One 8, e66235. 10.1371/journal.pone.0066235

Shobin, E., Bowley, M.P., Estrada, L.I., Heyworth, N.C., Orczykowski, M.E., Eldridge, S.A., Calderazzo, S.M., Mortazavi, F., Moore, T.L., Rosene, D.L., 2017. Microglia activation and phagocytosis: relationship with aging and cognitive impairment in the rhesus monkey. GeroScience 39, 199–220. 10.1007/s11357-017-9965-y

Wahl, D., Moreno, J.A., Santangelo, K.S., Zhang, Q., Afzali, M.F., Walsh, M.A., Musci, R.V., Cavalier, A.N., Hamilton, K.L., LaRocca, T.J., 2022. Nontransgenic Guinea Pig Strains Exhibit Hallmarks of Human Brain Aging and Alzheimer’s Disease. J. Gerontol. A. Biol. Sci. Med. Sci. 77, 1766– 1774. 10.1093/gerona/glac073

Welker, W., Johnson, J.I., Noe, A., 2010. Comparative mammalian brain collections: the domesticated guinea pig brain. Major Natl. Resour. Study Brain Anat. 2010.

Wixey, J.A., Lee, K.M., Miller, S.M., Goasdoue, K., Colditz, P.B., Tracey Bjorkman, S., Chand, K.K., 2019. Neuropathology in intrauterine growth restricted newborn piglets is associated with glial activation and proinflammatory status in the brain. J. Neuroinflammation 16, 5. 10.1186/s12974-018-1392-1

Zinni, M., Pansiot, J., Colella, M., Faivre, V., Delahaye-Duriez, A., Guillonneau, F., Bruce, J., Salnot, V., Mairesse, J., Knoop, M., Possovre, M.-L., Vaiman, D., Baud, O., 2021. Impact of Fetal Growth Restriction on the Neonatal Microglial Proteome in the Rat. Nutrients 13, 3719. 10.3390/nu13113719

